# Human-gut phages harbor sporulation genes

**DOI:** 10.1101/2023.01.19.524802

**Authors:** DA Schwartz, J Rodríguez-Ramos, M Shaffer, R Flynn, R Daly, KC Wrighton, Jay T. Lennon

## Abstract

Spore-forming bacteria are prevalent in mammalian guts and have implications for host health and nutrition. The production of dormant spores is thought to play an important role in the colonization, persistence, and transmission of these bacteria. Spore formation also modifies interactions among microorganisms such as infection by phages. Recent studies suggest that phages may counter dormancy-mediated defense through the expression of phage-encoded sporulation genes during infection, which can alter the transitions between active and inactive states. By mining genomes and gut-derived metagenomes, we identified sporulation genes that are preferentially encoded by phages that infect spore-forming bacteria. These included genes involved in chromosome partitioning, DNA damage repair, and cell wall-associated functions. In addition, phages contained homologs of sporulation-specific transcription factors, notably *spo0A*, the master regulator of sporulation, which could allow phages to control the complex genetic network responsible for spore development. Our findings suggest that phages could influence the formation of bacterial spores with implications for the health of the human gut microbiome, as well as bacterial communities in other environments.

**SIGNIFICANCE:** Phages acquire bacterial genes and use them to alter host metabolism in ways that enhance their fitness. To date, most auxiliary genes replace or modulate enzymes that are used by the host for nutrition or energy production. However, phage fitness is affected by all aspects of host physiology, including decisions that reduce metabolic activity of the cell. Here we focus on endosporulation, a complex and ancient form of dormancy found among the Bacillota that involves hundreds of genes. By coupling homology searches with host classification, we identify 31 phage-encoded homologs of sporulation genes that are mostly limited to phages infecting spore-forming bacteria. Nearly one-third the homologs recovered were regulatory genes suggesting that phages may manipulate host genetic networks by tapping into their control elements. Our findings also suggest a mechanism by which phages can overcome the defensive strategy of dormancy, which may be involved in coevolutionary dynamics of spore-forming bacteria.

## MAIN TEXT

Microbiomes in the human gut are made up of a diverse community of bacteria, archaea, and microeukaryotes, as well as well as viruses that infect these microorganisms (1). Members of the phylum Bacillota (formerly Firmicutes) include many spore-forming lineages such as *Bacillus* and *Clostridium*. These spore-forming members are dominant in healthy gut microbiomes, with many strains within this group also considered common intestinal pathogens (2, 3). Sporulation is a complex form of dormancy, involving hundreds of genes, that helps these bacteria contend with spatial and temporal variation in environmental conditions in human guts and facilitate transmission (2, 4).

Viruses of microbes, such as bacteriophages, play an important role in shaping gut microbiomes (1). Phage fitness is thought to be enhanced through the encoding of bacterial-like auxiliary metabolic genes (AMGs) that can reprogram and sustain host metabolism during infection (5). The acquisition of other, non-metabolic genes may allow phages to alter other aspects of bacterial physiology (6). One of the most important determinants of phage fitness is the metabolic activity of the host cell (7, 8). Bacterial metabolism spans orders of magnitude, ranging from exponential growth to being nearly inert when cells engage in certain types of dormancy such as sporulation (9, 10). By entering a state of reduced metabolic activity, microorganisms can defend themselves against phage attack (11, 12) altering selection in ways that could modify coevolutionary dynamics.

Previous work has demonstrated that some phage isolates encode sporulation genes (e.g. 13, 14, 15). In one example, homologs of sporulation-specific sigma factors (*sigG* and *sigF*) were identified in both lytic and lysogenic phages (13). These sigma factors are essential for the developmental transition from vegetative cell to and endospore (16). When expressed in a host (*Bacillus subtilis*), the phage-encoded sigma factors activate sporulation transcriptional pathways and depress spore yield by up to 99% (13). To date, there has not been any systematic analysis of the prevalence and distribution of sporulation genes in phages. Thus, it remains unknown whether modification of host sporulation is a common phage strategy. In this study, we search for homologs of sporulation genes in genomic and metagenomic data to determine whether phages employ this strategy in human gut microbiomes.

### Identifying sporulation homologs in viral genomes and metagenomes

We identified sporulation genes in viral genomes and metagenomic assembled genomes (vMAGS) using DRAM-v (17) (Fig. S1 and supplementary text). Specifically, we targeted homologs of well characterized sporulation genes found in *B. subtilis* and *Clostridioides difficile*. We reasoned that phage-encoded genes can only affect sporulation if they are in phages that infect a spore-forming host. We therefore designed an enrichment test to identify homologs of sporulation genes that were preferentially found in phages that infect spore-forming hosts. We first evaluated our search strategy by looking for sporulation genes in genomes of phage isolates, for which the host was known. Next, we applied the same approach to vMAGs assembled from human gut environments for which host predictions had been made (18, 19). To minimize potential for contamination by bacterial sequences, we inspected the annotations of 6,542 gut-derived vMAGs in which sporulation genes were detected, with an average of 117 vMAGs inspected per enriched sporulation gene (Fig. S2).

### Phages encode structural genes required for sporulation

Our search identified homologs of 31 phage-encoded homologs of sporulation genes (Table 1). These sporulation genes were enriched in phages that infect spore-forming hosts (Fig. 1). Many of the phage-encoded homologs were structural (i.e., non-regulatory) genes involved in an assortment of sporulation-related processes such as chromosome partitioning, DNA damage repair, and cell wall-associated functions (Table 1). The acquisition of these genes might allow phages to promote or impede specific steps of spore development, or its eventual germination. For example, phages may use chromosome segregation genes to increase the probability of entrapment (and survival) of the phage genome in the spore during the asymmetric division separating the developing spore from the mother cell (20). Alternatively, it is possible that some of these genes are used by phages for functions other than sporulation. Chromosome segregation genes are known to be used by phages that establish extrachromosomal, plasmid-like lysogeny (21). Likewise, cell wall hydrolases used by the host to restructure the cell during sporulation (*cwlJ*, *sleB*, *spoIID*) could be repurposed as endolysins to burst the host cell at the completion of the phage lytic cycle (22). Further experimental investigation will be required to establish the phage functions of the sporulation gene homologs that we have catalogued in this work.

**Table 1.** Sporulation genes detected in viral genomes and metagenomes. These genes were enriched in phages of spore-forming hosts and were validated to have a viral origin by manual inspection of annotations. KO represents the KEGG ortholog identifier. ‘Locus no.’ and `Gene` refer to the gene locus number and name of sporulation gene(s) associated with a KO. ‘BSU’ loci are from *Bacillus subtilis* (KEGG taxon T00010) and CD630 loci are from *Clostridioides difficile* (KEGG taxon T00487). Locus `Type` reflects whether a KO is a regulatory gene (R), structural gene (S), or a hypothetical of uncharacterized function (U). The `Function` column provides a description of the KO from Subtiwiki for *B. subtilis* or KEGG for *C. difficle*.

| KO | Locus no. | Gene | Type | Function |
| --- | --- | --- | --- | --- |
| K01356 | BSU_17850 | *lexA* | R | lexA repressor |
| K03086 | BSU_25200 | *sigA* | R | RNA polymerase sigma factor RpoD |
| K03091 | BSU_00980 | *sigH* | R | RNA polymerase sigma-H factor |
|  | BSU_15320 | *sigE* |  | RNA polymerase sigma-E factor |
|  | BSU_15330 | *sigG* |  | RNA polymerase sigma-G factor |
|  | BSU_23450 | *sigF* |  | RNA polymerase sigma-F factor |
|  | CD630_07720 | *sigF* | R | RNA polymerase sigma-F factor |
|  | CD630_12300 | *sigK* |  | RNA polymerase sigma-K factor |
|  | CD630_26420 | *sigG* |  | RNA polymerase sigma-G factor |
|  | CD630_26430 | *sigE* |  | RNA polymerase sigma-E factor |
| K04769 | BSU_560 | *spoVT* | R | stage V sporulation protein T |
|  | CD630_34990 | *spoVT* | R | Stage V sporulation protein T |
| K06283 | BSU_36420 | *spoIIID* | R | Stage III sporulation protein D |
|  | CD630_1260 | *spoIIID* | R | Stage III sporulation protein D |
| K06284 | BSU_370 | *abrB* | R | transition state regulatory protein AbrB |
| K07699 | BSU_24220 | *spo0A* | R | stage 0 sporulation protein A |
|  | CD630_12140 | *spo0A* | R | Stage 0 sporulation protein A |
| K07738 | CD630_26400 | *nrdR* | R | Transcriptional regulator, repressor NrdR family |
| K03496 | BSU_40970 | *parA* | R+S | sporulation initiation inhibitor protein Soj |
|  | CD630_36720 | *soj* | R+S | Transcriptional regulator, sporulation initiation inhibitor, chromosome partitioning protein |
| K00390 | BSU_10930 | *yitB* | S | phosphoadenosine phosphosulfate reductase |
| K00640 | CD630_15950 | *cysE* | S | Serine acetyltransferase (SAT) |
| K00820 | CD630_1200 | *glmS* | S | Glucosamine--fructose-6-phosphate aminotransferase [isomerizing] |
| K00974 | BSU_22450 | *cca* | S | CCA-adding enzyme |
| K01142 | BSU_40880 | *exoA* | S | Exodeoxyribonuclease, repair of oxidative DNA damage in spores |
| K01449 | BSU_02600 | *cwlJ* | S | cell wall hydrolase CwlJ |
|  | BSU_22930 | *sleB* |  | spore cortex-lytic enzyme |
|  | CD630_35630 | *NA* | S | putative spore cortex-lytic hydrolase |
| K02049 | BSU_30610 | *ytlC* | S | ABC transporter ATP-binding protein |
| K02343 | CD630_160 | *dnaX* | S | DNA polymerase III subunits gamma and tau |
| K03466 | BSU_16800 | *spoIIIE* | S | spore DNA translocase |
| K03497 | BSU_40960 | *parB* | S | stage 0 sporulation protein J |
|  | CD630_36710 | *spo0J* | S | Stage 0 sporulation protein J, site-specific DNA-binding protein |
| K03657 | CD630_7490 | *NA* | S | putative DNA helicase, UvrD/REP type |
| K03664 | BSU_33600 | *smpB* | S | SsrA-binding protein |
| K03698 | BSU_9930 | *yhaM* | S | 3'-5' exoribonuclease yhaM |
| K06381 | BSU_36750 | *spoIID* | S | stage II sporulation protein D |
|  | CD630_1240 | *spoIID* | S | Stage II sporulation protein D |
| K06412 | BSU_490 | *spoVG* | S | septation protein SpoVG |
|  | CD630_35160 | *spoVG* | S | Regulator required for spore cortex synthesis |
| K07171 | CD630_34610 | *EndoA* | S | Endoribonuclease toxin |
| K10716 | BSU_31322 | *yugO* | S | potassium channel protein YugO |
| K10979 | BSU_13410 | *ykoV* | S | DNA repair protein YkoV |
| K01448<sup>7</sup> | BSU_17410 | *cwlC* | S | mother cell lysis |
|  | BSU_01530 | *cwlD* |  | spore cortex peptidoglycan synthesis |
| K02647 | BSU_28670 | *ysfB* | U | hypothetical protein; similar to carbohydrate diacid transcriptional activator |
| K03469 | BSU_21970 | *ypeP* | U | hypothetical protein; similar to RNase HI |
| K07175 | BSU_14810 | *ylaK* | U | hypothetical protein; similar to PhoH |

**Fig. 1.**
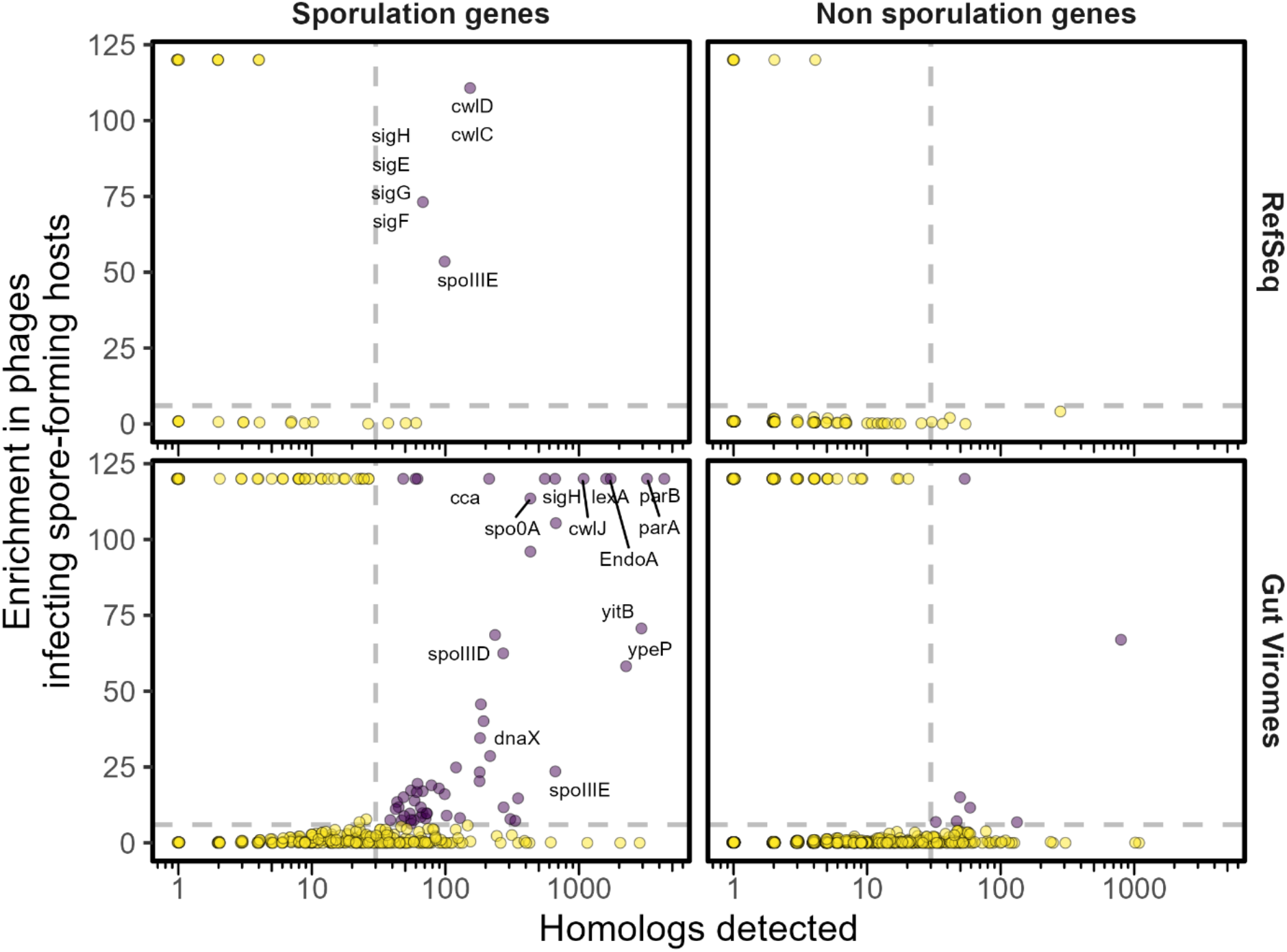
Sporulation genes enriched in phages that infect spore-forming bacteria. Homologs of sporulation and non-sporulation genes were identified in RefSeq isolate phage genomes and in published gut viromes (18, 19) using DRAM-v. A hypergeometric enrichment test was performed for each gene to evaluate if it was found in phages that infect spore-forming hosts more than the random expectation given the total number of phages. The enrichment result (*‒Log_10_ (hypergeometric P value)* on y-axis) is plotted as a function of the number of homologs detected (x-axis). Purple points represent enriched genes with an adjusted *P*-value < 10^−6^ (horizontal dashed line) and a sample size > 30 (vertical dashes line). Representative names of *B. subtilis* sporulation genes are provided for genes that were enriched and of viral origin.

### Phages also encode genes that regulate sporulation

Nearly one-third of the sporulation homologs (n = 9) identified in phage genomes and metagenomes are transcriptional regulators (Table 1). This finding is different than most examples of AMGs, where the phages control a metabolic process by phage-encoded enzymes, or by expression of modulators of host enzyme activity (5). It may be that phages manipulate host sporulation by interfering with the tightly regulated transcriptional program that is essential for this complex developmental process (2).

Such findings are consistent with recent experimental findings with sigma factors where the ectopic expression of phage-encoded *sigG* and *sigF* homologs altered the transcriptional program of *B. subtilis* resulting in reduced spore yield (13).

Most notable among phage-encoded regulators are homologs of *spo0A*, the master regulator of sporulation initiation that is conserved among all spore-forming bacteria (4). Interestingly, the homologs found in phages are truncated versions of *spo0A* that contain the DNA-binding effector domain, but not the receiver domain (Fig. S3). The latter is responsible for modifying the DNA-binding activity in response to environmental and physiological signals received via the phosphorelay signal-transduction system (23). The truncation suggests that phage-encoded *spo0A* may not require the normal host signals to activate or repress the initiation of host sporulation (24). Beside transcriptional regulators, phage genes included other potential post-transcriptional regulators (RNA binding *spoVG*, and translation-related genes *cca* and *smpB*). Taken together, bioinformatic findings here and laboratory results (13) suggest some phages may overcome dormancy defenses by targeting the regulation of sporulation. Compared to the use of structural genes, this is likely to be a more efficient strategy for altering the course of a complex cellular program.

### Phage-encoded sporulation genes occur in diverse environments

The recovery of phage-encoded sporulation genes is not restricted to the human gut. We identified sporulation genes in vMAGs originating from diverse environments (Table S1). Of the 30 sporulation genes identified in gut-derived vMAGs we found 23 that also occur in phages from terrestrial and aquatic environments (Fig S4). Thus, phage manipulation of sporulation may be a common phenomenon in environments where spore-forming bacteria are found.

### Implications and future directions

Sporulation is an ancient, complex, and important trait that contributes to the persistence and transmission of beneficial and pathogenic members of the mammalian gut microbiome. While sporulation can reduce virus infection, our analysis supports the view that phages may use host-like genes to overcome host defense mechanisms (25). Specifically, our study provides genomic and metagenomic evidence that phages encode multiple homologs of sporulation genes, which may influence the transition of bacteria between active and dormant states in host-associated and environmental ecosystems. The evolutionary drivers and ecological consequences of phage-encoded sporulation genes remain to be investigated (15). Our work demonstrates how partitioning phages by a specific host trait (e.g., sporulation) can be used to identify genes used by phages to influence the same host trait.

## CODE AND DATA AVAILABILITY

All code and data used in this study are available at https://github.com/LennonLab/spore_amg. In addition, prior to publication, all data and code will be made available on Zenodo.

## ACKNOWLEGEMENTS

Research was supported by the National Science Foundation (DEB-1934554 JTL and DAS, DBI-2022049 JTL, EAR-1847684 KCW), US Army Research Office Grant (W911NF-14-1-0411 JTL and W911NF-22-1-0014), the National Aeronautics and Space Administration (80NSSC20K0618 JTL).

