## Supplementary Materials for "Human-gut phages harbor sporulation genes"

### SUPPLEMENTARY TEXT

#### List of sporulation genes

To search for sporulation genes in phages, we compiled a list of sporulation-related genes for the two best studied endospore-forming bacteria: *Bacillus subtilis* and *Clostridioides difficile*. For *B. subtilis*, we identified 880 sporulation genes that have been experimentally evaluated and summarized in the Subtiwiki database (1) and other publications (2, 3) . For *C. difficile*, we identified a total of 350 sporulation genes. We included genes that were functionally annotated as being involved in sporulation (GenBank: CP016318.1) along with sporulation genes that were identified in at least two of four curated studies (2, 4-6). We mapped sporulation genes for both species to functional orthologous groups in the KEGG Orthology (KO) database using the KEGGREST R Package (7), which enabled detection of diverse sporulation gene homologs using hidden-Markov-model (HMM) based annotations by DRAM (see below).

#### Classification of host sporulation capacity

To evaluate if a phage's host is likely to be a spore-former, we compiled data on spore formation within families of the phylum Bacillota (formerly Firmicutes). Assuming that only members of the Bacillota form endospores, we assigned members of all other phyla as non-sporulators. Within the Bacillota we combined data on spore-formation from two previous studies (8, 9) . We classified families as spore-formers if >50% of the species identified by Browne et al. (9) were found to be spore formers. For 10 families whose spore-forming status differed between the two studies (8, 9), we assigned spore-forming status by manual inspection of the underlying data in these studies, supplemented by additional sources (10-12). Within 402 families from the Bacillota defined by the genome taxonomy database (GTDB v202), we assigned 165 as spore-formers and 92 as non-non-spore-formers, while 145 were not assigned due to insufficient data.

**Identification of vMAGs, manual curation of genomes, and detection of sporulation genes in phages** Publicly available datasets that were used in this study were downloaded from their respective repositories with their predicted host information (13, 14) and annotated using DRAM-v v1.3.1 to identify candidate homologs of sporulation genes (15). DRAM is a program that annotates genomes and assigns the genes to predefined metabolic modules. Using KO IDs corresponding to the curated list of sporulation genes, we supplemented DRAM with a sporulation module that is publicly available on the DRAM GitHub Repository (https://github.com/WrightonLabCSU/sporulation_utils). As previously described, DRAM's viral mode (DRAM-v) classifies metabolic genes in viral genomes as putative auxiliary metabolic genes (AMGs). It then provides a score of 1-5 that describes confidence metrics to annotation calls as well as distillate flags that aid in distinguishing between bacterial genes that originate from viral genomes and those from bacterial genome fragments erroneously classified as viral. For publicly available datasets that did not have available viral contigs, we downloaded the respective assemblies (Table S1) and then used VirSorter v1.0.3 with default settings to identify viral metagenome assembled genomes (vMAGS) from our metagenomics datasets (16). We selected high confidence vMAGs that were greater than or equal to 10kb in size and were assigned to categories 1-2 and 4-5 by VirSorter. These were then clustered into vMAGs at 95% ANI across 85% of the shortest contig using ClusterGenomes 5.1 (https://github.com/simroux/ClusterGenomes) according to established standards (17). We then used DRAM-v v1.3.1 with default settings for functional annotation.

To avoid misidentification of bacterial contaminants as bacterial genes encoded by viruses (i.e., AMGs), we supplemented our discovery pipeline with manual inspection and curation of vMAGs (18, 19). For each of the sporulation genes enriched in the analysis of gut viromes, we performed a manual inspection of large samples of annotated vMAGs containing those genes by plotting the genome maps with features of the DRAM annotations (Fig. S5). We only considered genes as good hits if they corresponded to high-confidence AMG categories 1-3 as suggested by DRAM-v default settings (15). Briefly, putative virus-encoded sporulation genes were considered truly viral if they were nested between viral hallmark genes (annotations containing the words "virion", "capsid", "tail", "terminase", "baseplate", "phage", "virus", "reverse transcriptase" or "head") and genes with no annotation (including hypothetical genes and genes of unknown function). We also excluded sporulation genes that were at or near the edge of a scaffold (DRAM AMG flag "F"). Finally, we excluded sporulation genes that were found in regions having characteristics of bacterial origin such as regions in which most genes were annotated as non-viral by KEGG or PFAM in the DRAM annotations, regions being indicative of bacterial transposons with no viral annotations (DRAM AMG Flag "T"), or regions having many gene direction switches. Sporulation genes that were detected >10 times in clearly viral contig regions are considered likely viral sporulation genes.

*Enrichment of sporulation genes in viral isolates* – We reasoned that phage-encoded genes can only affect sporulation if they are encoded by phages that infect a host capable of forming spores. We first searched for sporulation genes in phage genomes from the RefSeq viral database (v202). This is a database curated and annotated by the National Center for Biotechnology Information (NCBI) with many genomes of phage isolates, so there is little concern of bacterial contamination in the sequences, and the identity of the hosts is known. We conducted the proof-of-concept test using only *B. subtilis* sporulation genes. Phage hosts identified using the virus-host database (20) were classified as spore-formers or non-spore-formers based on family-level taxonomy, as described above. We were able to classify host sporulation capacity of 3,650 phages, of which 257 are known to infect spore-forming hosts. For each candidate gene, we conducted a hypergeometric enrichment test to determine if the gene was detected in phages infecting spore-formers more than expected by a random draw of phages from the pool of 3,650 phages. *P*-values were adjusted for multiple comparisons using the Benjamini-Hochberg method (21) . Sporulation genes with less than 30 observations were excluded from our analysis.

*Enrichment of sporulation genes in viral metagenomes by host sporulation* – To expand our search of viral sporulation genes beyond phage isolates, we applied the same enrichment approach described above using two previously published datasets of vMAGs assembled from human gut samples. Typically, there is a high relative abundance of spore-forming Bacillota in these environments, which was confirmed by applying our spore-forming classification to host predictions that were made as part of the original studies (13, 14). We annotated the vMAGs from these two datasets using DRAM-v equipped with the full sporulation module based on sporulation genes from both *B. subtilis* and *C. difficile*. We matched the available host predictions (published alongside these datasets) to our list of sporulators at the family level. We classified host sporulation capacity of 53,624 vMAGs, of which 25,630 are predicted to infect spore-forming hosts. Enrichment tests for sporulation genes found in viral contigs of both datasets were conducted as described above. In addition to the human gut samples, we used DRAM-v equipped with the full sporulation module to annotate vMAGs from a variety of metagenomic datasets originating from aquatic, terrestrial and host associated environments (Table S1).

**Multiple sequence alignment of viral and bacterial *spo0A* genes** – To compare viral and bacterial homologs of spo0A we retrieved bacterial *spo0A* protein sequences from the Clusters of Orthologous Genes (COG) database (COG5801 in v2020 (22). We clustered the sequences using CD-HIT-est (23) at 65% identity over 90% of the largest sequence (-c 0.65 -aS 0.9). Bacterial sequences were aligned together with sequences from virome scaffolds that passed manual inspection (see above) using MUSCLE v5 (24). Multiple sequence alignments were visualized using the ggmsa (25) package in R (26).

**Code and data availability** – All code and data used in the analyses in this study are available at github.com/LennonLab/spore_amg as well as https://github.com/jrr-microbio/sporulation_project. In addition, prior to publication, all data and code will be made available on Zenodo.

### SUPPLEMENTARY FIGURES

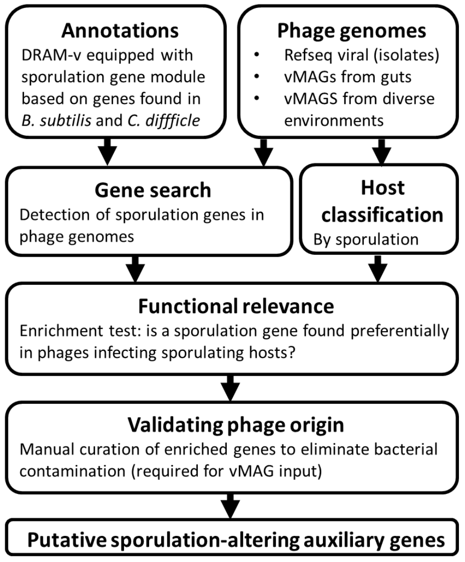


**Fig. S1.** Workflow for identification of phage homologs of sporulation genes. vMAG: viral metagenome-assembled genome.

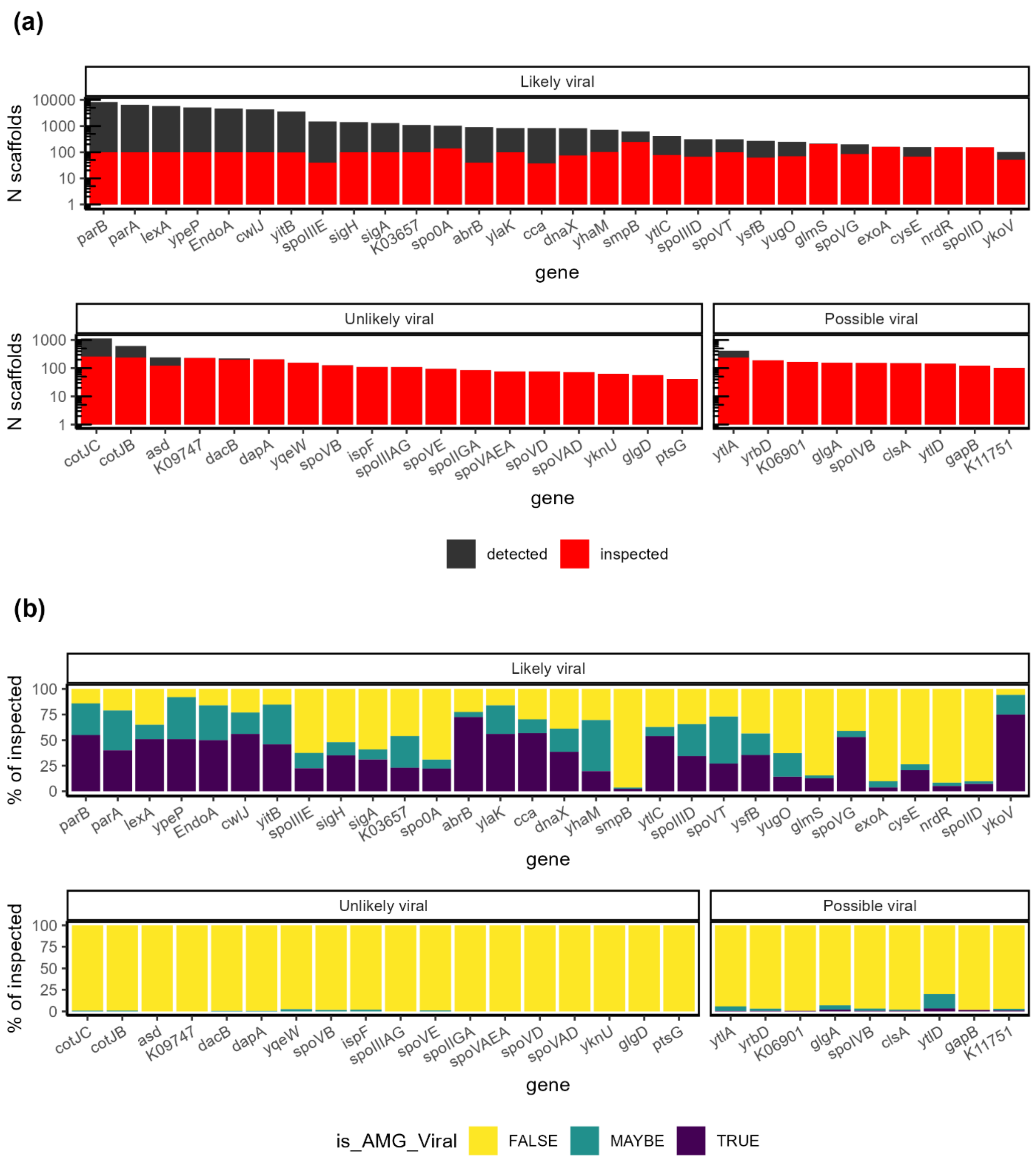


**Fig. S2.** Manual inspection of gut-derived viral metagenomic assembled genomes (i.e., scaffolds) containing sporulation genes from (13, 14). In each panel the data are separated between genes for which a few examples of true viral origin were observed ("Likely viral"), never observed ("Unlikely viral") or observed in very few instance (<5 scaffolds; "Possible viral"). **(a)** The number of scaffolds in which each of the sporulation genes was detected, and the number of scaffolds manually inspected. **(b)** The fraction of the inspected scaffolds in which the focal sporulation gene is likely of viral origin ("TRUE"), likely originates from bacterial contamination ("FALSE"), or difficult to determine ("MAYBE").

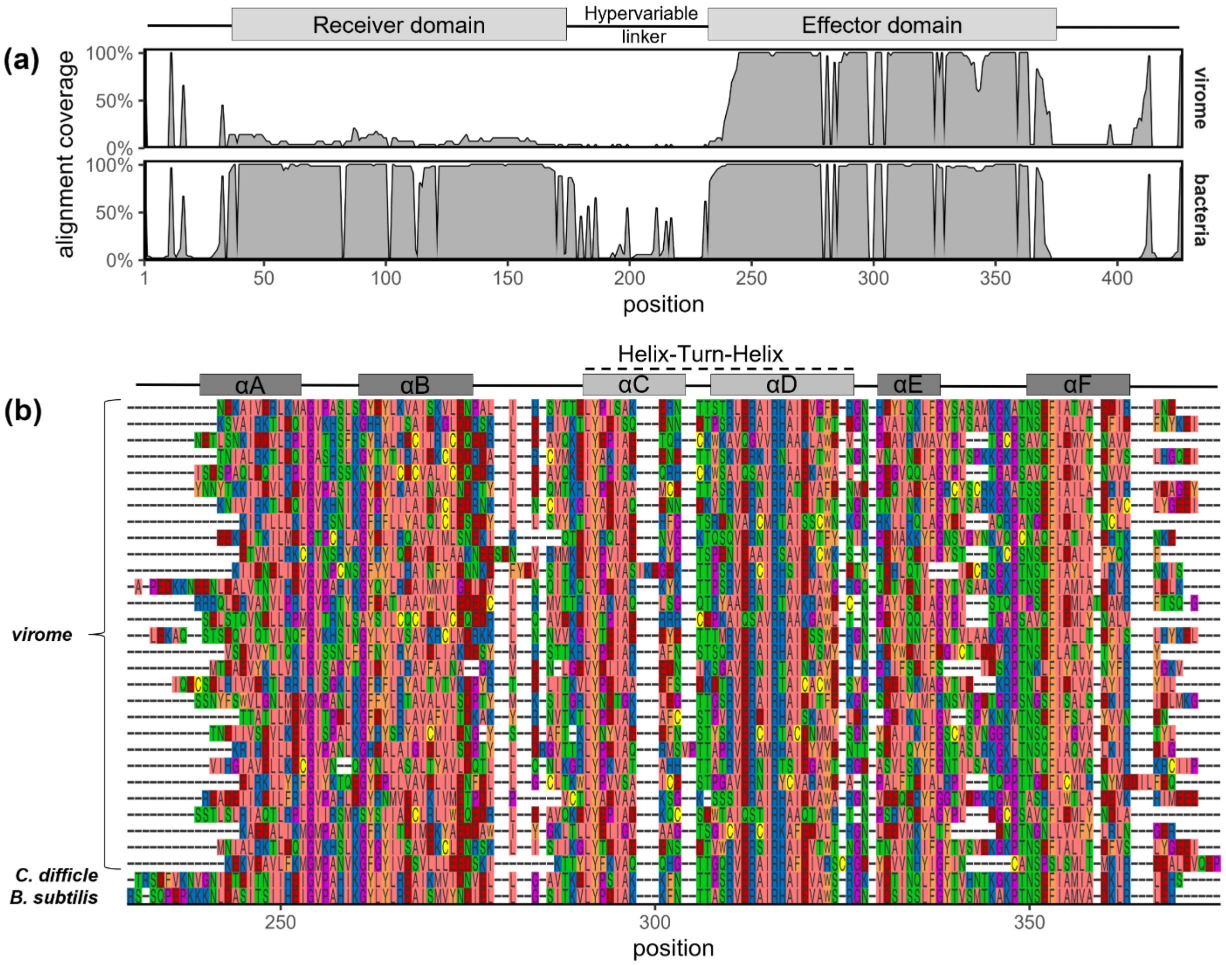


**Fig. S3**. Multiple sequence alignment of protein sequences of *spo0A* homologs from viromes (n = 29) and diverse bacteria (n = 57; from COG5801). **(a)** summary of alignment coverage showing the percent of non-gap characters at each position, separated by the source of the sequence. *Spo0A* from viromes align to the C-terminal effector domain of the bacterial genes. **(b)** focus on effector domain, showing all virome sequences aligned with *spo0A* of model spore-forming bacteria *Clostridioides difficile* and *Bacillus subtilis*. The alpha helices of the effector domain are indicated above the sequences. Sequence colors correspond to physicochemical properties of amino acids, using the Zappo coloring scheme (1). Information on functional and structural domains from (2). Plot made using ggmsa (3).


1. https://www.jalview.org/help/html/colourSchemes/zappo.html
2. Lewis, R.J., et al., The trans-activation domain of the sporulation response regulator Spo0A revealed by X-ray crystallography. Molecular Microbiology, 2000. **38**(2): p. 198-212.
3. Zhou, L., et al., *ggmsa: a visual exploration tool for multiple sequence alignment and associated data.* Briefings in Bioinformatics, 2022.

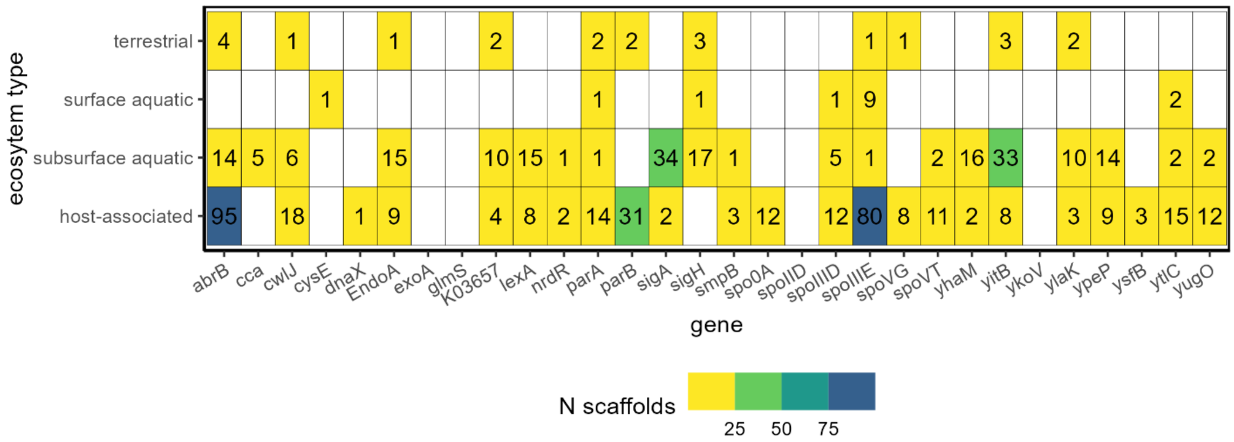


**Figure S4.** Sporulation genes detected in viral metagenomic assembled genomes (=scaffolds) from diverse ecosystems. “Host-associated” ecosystem includes samples derived from human guts, but these are different from the datasets discussed in main results. The number of scaffolds in which a gene was detected by DRAM-v and validated by manual inspection of scaffold annotations is shown. Empty cells indicate no scaffolds detected. See supplementary text for details on methods and samples.

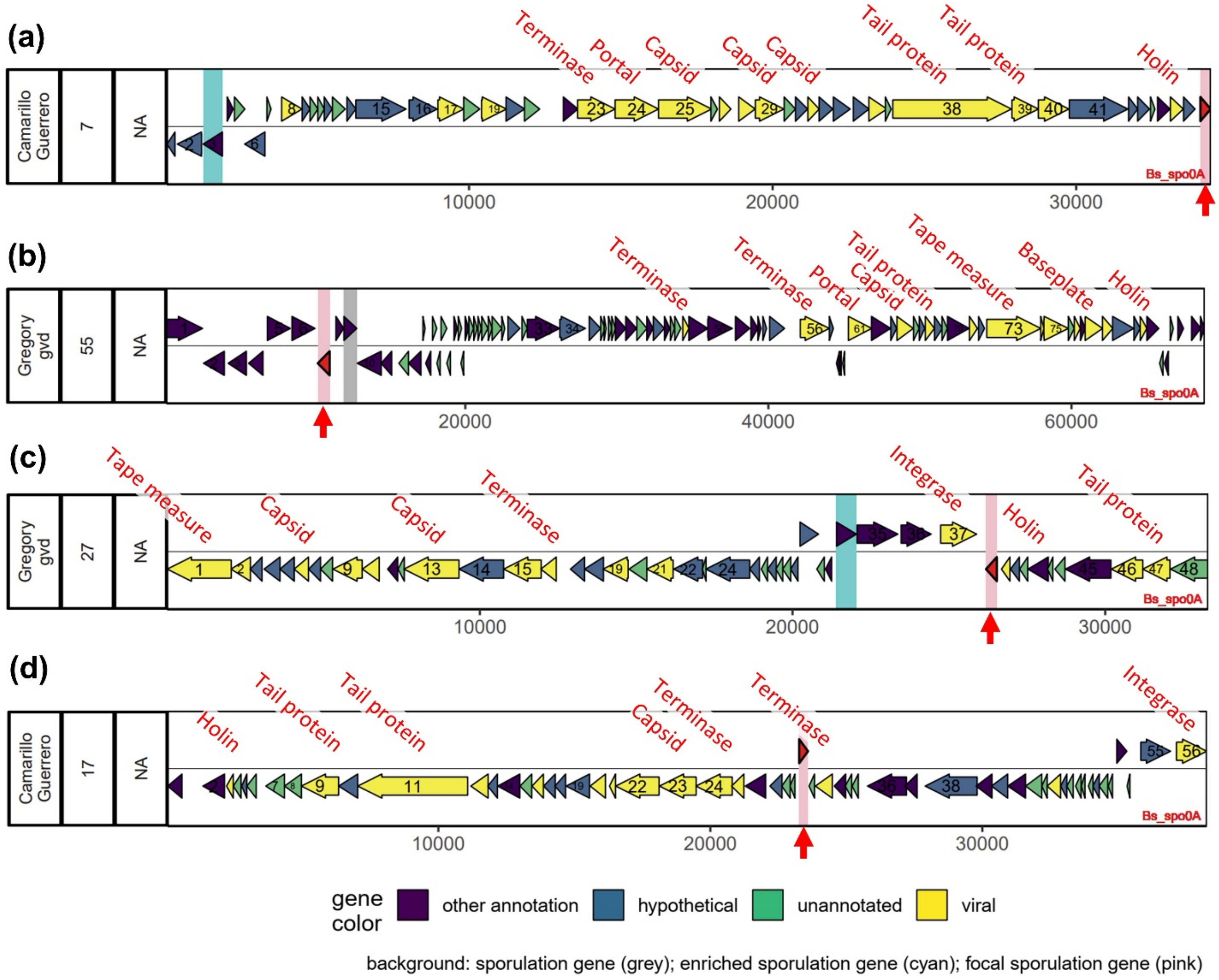


**Figure S5**. Examples of color-coded genome maps used in manual curation. Arrows correspond to gene annotations along scaffolds generated by DRAM-v. Gene direction is indicated by arrow direction and vertical position. Gene colors summarize functional annotations of individual genes from KEGG and PFAM. Annotations containing the terms "virion", "capsid", "tail", "terminase", "baseplate", "phage", "virus", "reverse transcriptase" or "head" were classified as hallmark viral genes. Genes having an annotation that was not viral or hypothetical were classified as “other annotation. The sporulation gene of interest (*spo0A* in these examples) is marked by a red arrow with a pink background, while other sporulation genes are marked with cyan or grey background, indicating genes enriched in phages of sporulating hosts, or not, respectively. **(a)** Rejected, as the *spo0A* gene is found at the edge of the scaffold. **(b)** Rejected, scaffold region with *spo0A* has many non-viral annotated genes and a high frequency of changes in gene direction, both indicating this is a bacterial genome fragment. The right-hand side of the scaffold appears to be viral based on the presence of multiple hallmark viral genes. **(c-d)** Accepted, in both case *spo0A* is nested between hallmark viral genes in scaffolds that otherwise contain many genes lacking annotation and few switches of gene direction. As needed the raw annotation files for a scaffold were also inspected. The viral gene annotations were added manually in these plots to illustrate that. Scaffold labels on the left indicate dataset name, scaffold index within dataset (indexed separately for each gene), and virsorter category (NA, not available for these scaffolds).

**Human-gut phages harbor sporulation genes**

### SUPPLEMENTARY TABLES

Table S1. Viromes analyzed in this study

| Set name | Ecosystem type | Description | Source |
|---|---|---|---|
| Gut Virome Database | host-associated | human gut study | (1) |
| Gut Phage Database | host-associated | human gut study | (2) |
| Anaerobic_Digestors | host-associated | incubations using sludge from wastewater treatment plant in Ft Collins, CO, digesting food waste from CSU | Wrighton, Unpublished |
| ARDEC_agriculture | terrestrial | soil samples from CSU's Agricultural Research, Development and Education Center (ARDEC) under a variety of cover crops | Wrighton, Unpublished |
| Columbia_River | surface aquatic | sediment samples from Columbia River, WA | (3) |
| CT_feces | host-associated | incubation of human feces (from methylamines study) amended with purified condensed tannin | Wrighton, unpublished |
| CT_soil | terrestrial | incubation of wetland soil (from OWC study) amended with purified condensed tannin | (4) |
| East_River | surface aquatic | sediment and pore water samples from East River, CO | (5) |
| Fire_soils | terrestrial | soil samples across a wildfire burn grandient in northern CO and southern WY | (6) |
| Frac_DJ_basin | subsurface aquatic | fracking wells in Colorado (unpublished data) | Wrighton, Unpublished |
| Frack_STACK | subsurface aquatic | fracking wells in Oklahoma | (7) |
| Fractured_shale | subsurface aquatic | Fracking wells in Appalachian Basin | (8) |
| Human_Microbiome_Project | host-associated | human gut study | (9) (*these were described in this paper) |
| Huttenhower | host-associated | human gut study, reads were from this dataset were assembled (unpublished) | (10) |
| Lake_Yojoa | surface aquatic | water column samples from Lake Yojoa, Honduras | Wrighton, Unpublished |
| Methylamines | host-associated | human fecal samples from healthy adults (from Cincinnati OH) | (11) |
| Moose_rumen | host-associated | rumen fluid samples from a moose in Alaska | (12) |

| | | | |
|---|---|---|---|
| Ohio_groundwater | subsurface aquatic | groundwater samples from 3 sites in OH | (13) |
| OWC_wetlands | surface aquatic | samples from wetland on shore of Lake Erie, CO | (14) |
| Prado_wetlands | surface aquatic | water samples from a constructed wetland in CA | (15) |
| Prairie_potholes | surface aquatic | sediment samples from Prairie Pothole region (lakes) in SD | (16) |
| Salmonella_chow | host-associated | feces from mice with and without Salmonella infection (fed normal mouse chow) | (17) |
| Salmonella_HFD | host-associated | feces from mice with and without Salmonella infection (fed a High Fat Diet) | Wrighton, Unpublished |
| Stordalen_mire | terrestrial | soil samples across a permafrost thaw gradient from Stordaline Mire Sweden (published data, viral-centric paper) | (18) |


1. Gregory AC, Zablocki O, Zayed AA, Howell A, Bolduc B, Sullivan MB. 2020. The gut virome database reveals age-dependent patterns of virome diversity in the human gut. Cell host & microbe 28:724-740. e8.
2. Camarillo-Guerrero LF, Almeida A, Rangel-Pineros G, Finn RD, Lawley TD. 2021. Massive expansion of human gut bacteriophage diversity. Cell 184:1098-1109. e9.
3. Rodríguez-Ramos JA, Borton MA, McGivern BB, Smith GJ, Solden LM, Shaffer M, Daly RA, Purvine SO, Nicora CD, Eder EK. 2022. Genome-resolved metaproteomics decodes the microbial and viral contributions to coupled carbon and nitrogen cycling in river sediments. bioRxiv.
4. McGivern BB, Tfaily MM, Borton MA, Kosina SM, Daly RA, Nicora CD, Purvine SO, Wong AR, Lipton MS, Hoyt DW. 2021. Decrypting bacterial polyphenol metabolism in an anoxic wetland soil. Nature communications 12:1-16.
5. Saup C, Bryant S, Nelson A, Harris K, Sawyer A, Christensen J, Tfaily M, Williams K, Wilkins M. 2019. Hyporheic zone microbiome assembly is linked to dynamic water mixing patterns in snowmelt-dominated headwater catchments. Journal of Geophysical Research: Biogeosciences 124:3269-3280.
6. Nelson AR, Narrowe AB, Rhoades CC, Fegel TS, Daly RA, Roth HK, Chu RK, Amundson KK, Young RB, Steindorff AS. 2022. Wildfire-dependent changes in soil microbiome diversity and function. Nature microbiology 7:1419-1430.
7. Amundson KK, Borton MA, Daly RA, Hoyt DW, Wong A, Eder E, Moore J, Wunch K, Wrighton KC, Wilkins MJ. 2022. Microbial colonization and persistence in deep fractured shales is guided by metabolic exchanges and viral predation. Microbiome 10:1-15.
8. Daly RA, Roux S, Borton MA, Morgan DM, Johnston MD, Booker AE, Hoyt DW, Meulia T, Wolfe RA, Hanson AJ. 2019. Viruses control dominant bacteria colonizing the terrestrial deep biosphere after hydraulic fracturing. Nature microbiology 4:352-361.
9. Shaffer M, Borton MA, McGivern BB, Zayed AA, La Rosa SL, Solden LM, Liu P, Narrowe AB, Rodríguez-Ramos J, Bolduc B. 2020. DRAM for distilling microbial

metabolism to automate the curation of microbiome function. Nucleic acids research 48:8883-8900.
10. Lloyd-Price J, Arze C, Ananthakrishnan AN, Schirmer M, Avila-Pacheco J, Poon TW, Andrews E, Ajami NJ, Bonham KS, Brislawn CJ. 2019. Multi-omics of the gut microbial ecosystem in inflammatory bowel diseases. Nature 569:655-662.
11. Borton MA, Shaffer M, Hoyt DW, Jiang R, Ellenbogen J, Purvine S, Nicora CD, Eder EK, Wong AR, Smulian AG, Lipton MS, Krzycki JA, Wrighton KC. 2022. Targeted curation of the gut microbial gene content modulating human cardiovascular disease. bioRxiv doi:10.1101/2022.06.20.496735:2022.06.20.496735.
12. Solden LM, Naas AE, Roux S, Daly RA, Collins WB, Nicora CD, Purvine SO, Hoyt DW, Schückel J, Jørgensen B. 2018. Interspecies cross-feeding orchestrates carbon degradation in the rumen ecosystem. Nature microbiology 3:1274-1284.
13. Danczak R, Johnston M, Kenah C, Slattery M, Wrighton KC, Wilkins MJ. 2017. Members of the Candidate Phyla Radiation are functionally differentiated by carbon-and nitrogen-cycling capabilities. Microbiome 5:1-14.
14. Angle JC, Morin TH, Solden LM, Narrowe AB, Smith GJ, Borton MA, Rey-Sanchez C, Daly RA, Mirfenderesgi G, Hoyt DW. 2017. Methanogenesis in oxygenated soils is a substantial fraction of wetland methane emissions. Nature communications 8:1-9.
15. Reilly JF, Horne AJ, Miller CD. 1999. Nitrate removal from a drinking water supply with large free-surface constructed wetlands prior to groundwater recharge. Ecological Engineering 14:33-47.
16. Dalcin Martins P, Danczak RE, Roux S, Frank J, Borton MA, Wolfe RA, Burris MN, Wilkins MJ. 2018. Viral and metabolic controls on high rates of microbial sulfur and carbon cycling in wetland ecosystems. Microbiome 6:1-17.
17. Leleiwi I, Rodriguez-Ramos J, Shaffer M, Sabag-Daigle A, Kokkinias K, Flynn RM, Daly RA, Kop LF, Solden LM, Ahmer BMM, Borton MA, Wrighton KC. 2022. Exposing New Taxonomic Variation with Inflammation – A Murine Model-Specific Genome Database for Gut Microbiome Researchers. bioRxiv doi:10.1101/2022.10.24.513540:2022.10.24.513540.
18. Emerson JB, Roux S, Brum JR, Bolduc B, Woodcroft BJ, Jang HB, Singleton CM, Solden LM, Naas AE, Boyd JA. 2018. Host-linked soil viral ecology along a permafrost thaw gradient. Nature microbiology 3:870-880.
